# Chemical inhibition of a CBASS anti-bacteriophage defense

**DOI:** 10.1101/2025.09.19.677395

**Authors:** Chengqian Zhang, Olga Rechkoblit, Joseph P. Gerdt

## Abstract

Multidrug-resistant (MDR) bacteria pose a significant public health challenge, underscoring the urgent need for innovative antibacterial strategies. Bacteriophages (phages), viruses that specifically target bacteria, offer a promising alternative; however, bacterial immune defenses often limit their effectiveness. Developing small-molecule inhibitors of these defenses can facilitate mechanistic studies and serve as adjuvants to enhance phage therapy. Here, we identify novel inhibitors targeting the bacterial cyclic oligonucleotide-based anti-phage signaling system (CBASS) effector Cap5. Cap5 is an HNH endonuclease activated by a cyclic nucleotide to degrade genomic DNA in virally infected cells, leading to cell death through abortive infection. Guided by the crystal structure of the Cap5 SAVED domain bound to its activating ligand, we performed structure-guided virtual screening to identify candidate inhibitors. Biochemical assays revealed that approximately 16% of the top docking hits exhibited inhibitory activity. Further cellular assays demonstrated that one potent compound could enter *E. coli* cells and inhibit Cap5 activity. Our integrated approach—combining structure-based virtual screening with biochemical validation—provides a robust framework for discovering small-molecule inhibitors of bacterial immune defenses to advance adjunctive therapies and deepen our understanding of phage-bacteria interactions.

## INTRODUCTION

Bacteria and their viruses, bacteriophages, have engaged in an evolutionary arms race for billions of years.^1^ Phages infect and kill a significant proportion of the bacteria on Earth daily,^2^ making them key players in shaping microbiomes and valuable tools in eradicating problematic bacteria.^3^ With the rising prevalence of antibiotic resistance, bacteriophages may become essential treatment options for multidrug-resistant (MDR) bacterial infections.^4^ To defend against phage attack, bacteria have evolved diverse defense immune systems.^5–7^ Some of the most common defenses have found extensive biotechnological applications (e.g., restriction enzymes^8^ and CRISPR^9, 10^). Beyond these ‘classic’ immune systems, hundreds of others have been discovered in the past few years.^11–16^ Many of these recently discovered immune systems are only partially characterized at a biochemical and functional level.

We believe that chemical inhibitors will prove to be valuable tools in the further study of these anti-phage immune systems.^17–19^ Furthermore, selective inhibition of individual immune systems may reveal the significance of each system for resistance to different families of phages. Finally, inhibitors may pave the way for combination therapies where phages and inhibitors work synergistically to enhance the efficacy of bacteriophage-based treatments against phage-resistant bacteria.

Many of the recently discovered immune systems function through a mechanism termed ‘abortive infection’, whereby infected bacterial cells undergo cell death to hinder phage propagation.^20^ Of these, cyclic-oligonucleotide-based anti-phage signaling systems (CBASS) are among the most widespread.^15, 21–24^ Upon infection, phage components activate the CBASS synthase enzyme (a cGAS/DncV-like nucleotidyltransferase, CD-NTase) to produce a cyclic nucleotide second messenger. The cyclic nucleotide then binds and activates the CBASS effector protein to kill an infected bacteria cell thus preventing the spread of phage particles within the bacterial cell population.^23–25^ The CBASS systems encode a diverse set of effectors to perform the cell suicide function ranging from degradation of bacterial genomic DNA,^25–27^ depletion of essential cellular metabolites^28, 29^ or damage to the bacterial membrane.^21, 30^

In this work, we focus on the CBASS Cap5 effector activated by 3′,2′-cGAMP (hereafter abbreviated as cGAMP in this manuscript) (Fig. 1A).^31–33^ The Cap5 protein contains an HNH DNA endonuclease domain coupled to a SAVED domain^25^ that binds the activating cyclic nucleotide ligand. Upon binding the ligand, Cap5 assembles into a tetramer composed of two crisscrossed dimers.^32^ In this configuration, two HNH domains—one in each dimer—are in a catalytically active state, ready to nonspecifically cleave both strands of bacterial DNA.^33^ The cyclic dinucleotide binds between the SAVED domains of each crisscross dimer. Notably, the ligand-binding pocket is primarily formed by the SAVED domain of the HNH-activated protomer within the crisscross dimer, while the SAVED domain of the second, HNH-inactive protomer forms a ‘lid’ over the pocket.

Here, to discover the inhibitors of a CBASS system, we virtually screened a library of commercially available compounds for binding to the single SAVED domain of *Lactococcus lactis* Cap5,^31^ which forms the primary cGAMP binding pocket.^32^ We then verified that six out of the 37 top ‘hits’ from the virtual screen are indeed capable of inhibiting cGAMP-activated nuclease activity of *L. lactis* Cap5 (*Ll*Cap5). Furthermore, we show that the homologous Cap5 from the plant pathogen *Pseudomonas syringae* (*Ps*Cap5) was inhibited by three of these six *Ll*Cap5 inhibitors. Additionally, a fourth compound—also identified through *in silico* screening—selectively inhibited *Ps*Cap5 but failed to inhibit *LlCap5.* Finally, we found that one of our inhibitors permeated into *Escherichia coli* cells and could inhibit *Ps*Cap5 within live cells. This finding establishes the efficacy of the strategy to chemically resensitize CBASS-containing bacteria (including pathogens like *P. syringae*) to bacteriophages.

## RESULTS

### Virtual screening prioritizes several potential inhibitors of the *Ll*Cap5 nuclease

We hypothesized that it is possible to identify compounds that bind the primary cGAMP-binding site of the SAVED domain, thereby competitively inhibiting Cap5 activation. To design such inhibitors, we used the virtual screening feature of CCDC (Cambridge Crystallographic Data Centre) GOLD^34^ to prioritize commercially available compounds predicted to bind the cGAMP site of the *Ll*Cap5 SAVED domain, which was co-crystallized with 3′,2′-cGAMP (PDB ID: 7RWS,^31^ Fig. 1B). To validate the accuracy of our docking strategy, we first re-docked the 3′,2′-cGAMP ligand into its binding pocket to ensure that the software placed it in the native configuration. The re-docked structure closely matched the original crystal structure with a calculated RMSD for the ligand of 0.55 Å, confirming the reliability of our approach.^35^ With confidence in our docking protocol, we then proceeded with a high-throughput virtual screen. An initial screen was conducted on a curated library of 8 million ‘lead-like’ compounds utilizing the CCDC GOLD supercomputer cluster’s virtual screening function. The top 70,000 compounds, based on their Chemical Piecewise Linear Potential (Chem-PLP)^36^ score, were then subjected to a higher accuracy docking screen. Following the refined screening, we collected the 102 highest-scoring compounds. After excluding compounds that were not readily available for purchase, we roughly binned the remaining 64 compounds into three groups (A, B, and C) by structural similarity using chemical fingerprinting,^37^ principal component analysis, and K-means analysis (Fig. 1E–F, Fig. S1–S3).^38^

**Figure 1.**
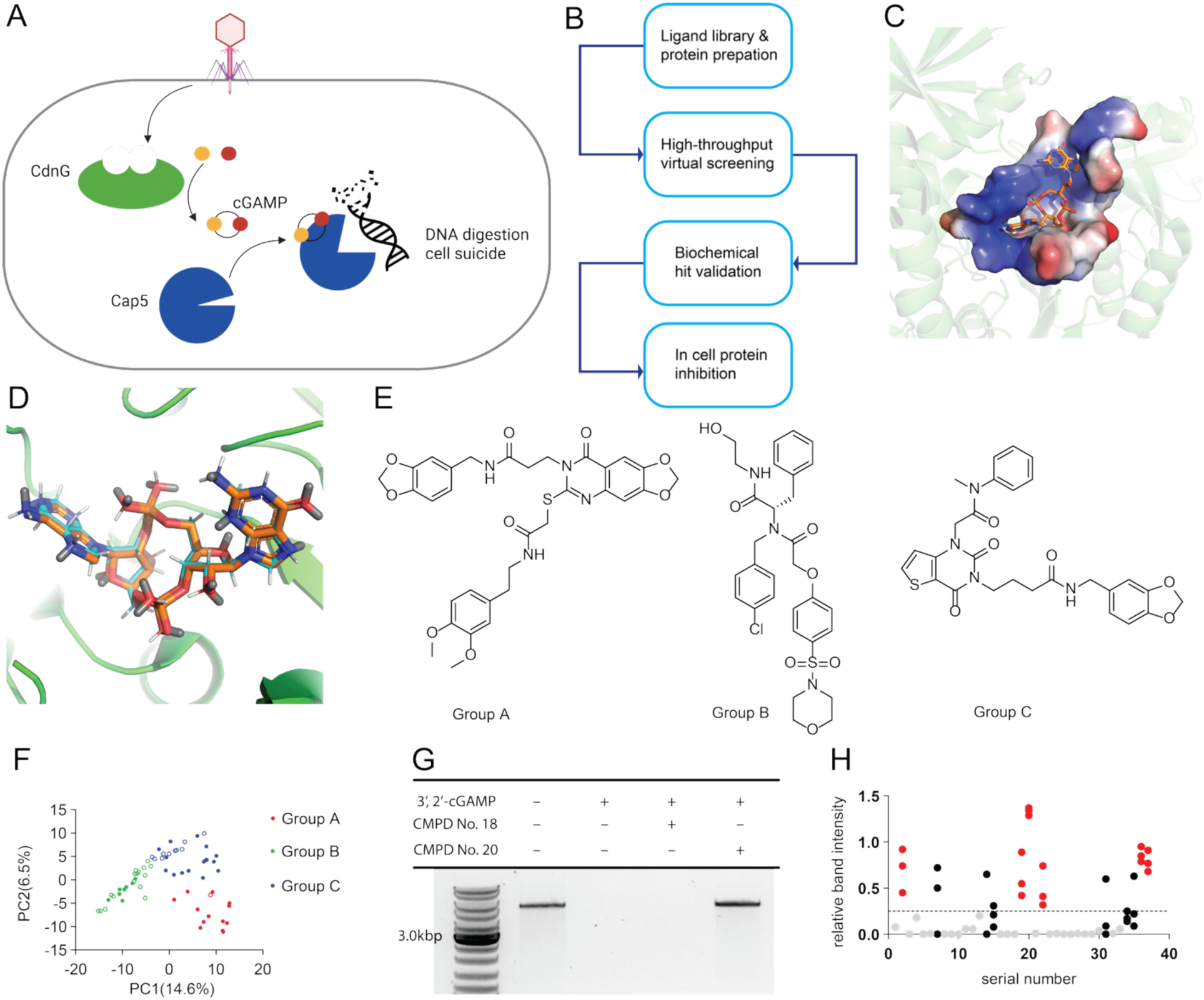
Virtual screening and biochemical validation. A) Schematic of the protection mechanism of the CBASS defense system with a Cap5 effector. Upon infection by a phage, the CdnG cyclase enzyme synthesizes a cyclic dinucleotide that binds and activates Cap5 effector protein triggering the death of the infected cell. B) Overview of the virtual screening scheme using GOLD.^34^ C) Crystal structure of the cGAMP activating ligand bound to the SAVED domain of the *L. lactis* Cap5 endonuclease (PDB ID: 7RWS).^31^ The pocket is shown as an electrostatic potential surface, with blue indicating positive charge, and red indicating negative charge. D) Re-docking of the cGAMP ligand, showing an alignment between the redocked and the experimental crystal structure. The crystal structure of 3′,2′-cGAMP bound to *Li*Cap5 SAVED is shown in orange, while the re-docked structure is shown in cyan (generated from early termination when the top three solutions are within 1 Å RMSD of each other). E) Representative structures of the three compound groups identified through K-means clustering on PCA from chemical fingerprints. Representative structures were chosen by medoid-based representative selection. For all structures, see Figures S1-3. F) PCA (Principal Component Analysis) of RDKit Daylight-like fingerprints of the top 64 commercially available compounds by Conda.^39^ Filled circles represent the compounds that were selected for biochemical testing. Empty circles represent compounds that were not tested biochemically. G) Example of an agarose gel for evaluation of DNA degradation activity of Cap5 endonuclease. Cap5 endonuclease activated by cGAMP ligand fully digests the DNA plasmid (band at 4811 kbps), whereas a DNA band can be observed and quantified when Cap5 is inactive/inhibited. H) Biochemical screening results for inhibition of *Ll*Cap5 nuclease. Band intensities were normalized to an undigested control band, with a value of one indicating a completely undigested band. Grey dots represent compounds that were inactive in the first test. Black dots show compounds that initially appeared active, but replicate experiments proved their activity to be inconsistent (at least one dot is below threshold). Red dots show compounds that repeatedly protected DNA from digestion by *Ll*Cap5. The threshold for activity was set at 0.4 (i.e., slightly less than half of the DNA band intensity remaining).

### Six Cap5 inhibitors were validated to be biochemically active

Among the 64 compounds, 37 representatives spanning the groups A, B, and C as well as overall chemical space, were purchased for biochemical validation (Fig. 1F and Fig. S1-S3). We anticipated that effective *Ll*Cap5 inhibitors would reduce the DNA cleavage activity of *Ll*Cap5 in the presence of its activating ligand (3′,2′-cGAMP), thereby preserving a detectable band of plasmid DNA on an agarose gel (Fig. 1G). Of the 37 compounds we tested, 27 failed to inhibit *Ll*Cap5 in the initial screen (Fig. 1H, grey dots). 10 others appeared to have some inhibitory activity, maintaining at least 40% of the undigested DNA band intensity. Upon retesting, six reproducibly inhibited *Ll*Cap5 (Fig. 1H, each replicate is a separate red circle), but four failed to replicate *Ll*Cap5 inhibition (Fig. 1H, each replicate is a separate black circle). Due to this inconsistency, these four compounds were excluded from further investigation. The six validated inhibitors display diverse chemical structures (Fig. 2A). Inhibition assays performed over a range of inhibitor concentrations revealed that most inhibitors have IC_50_s above 100 µM. The most potent compounds were **20** and **22**, with IC_50_s of 120 ± 60 μM and 58 ± 16 μM, respectively (Fig. 2B). The validated inhibitors represented both structural groups B (**19, 22, 37**) & C (**2, 20, 36**). Notably, many hits contain pyrimidone and purine-like substructures that mimic the nucleobases of the native ligand, likely contributing to their binding affinity.

**Figure 2.**
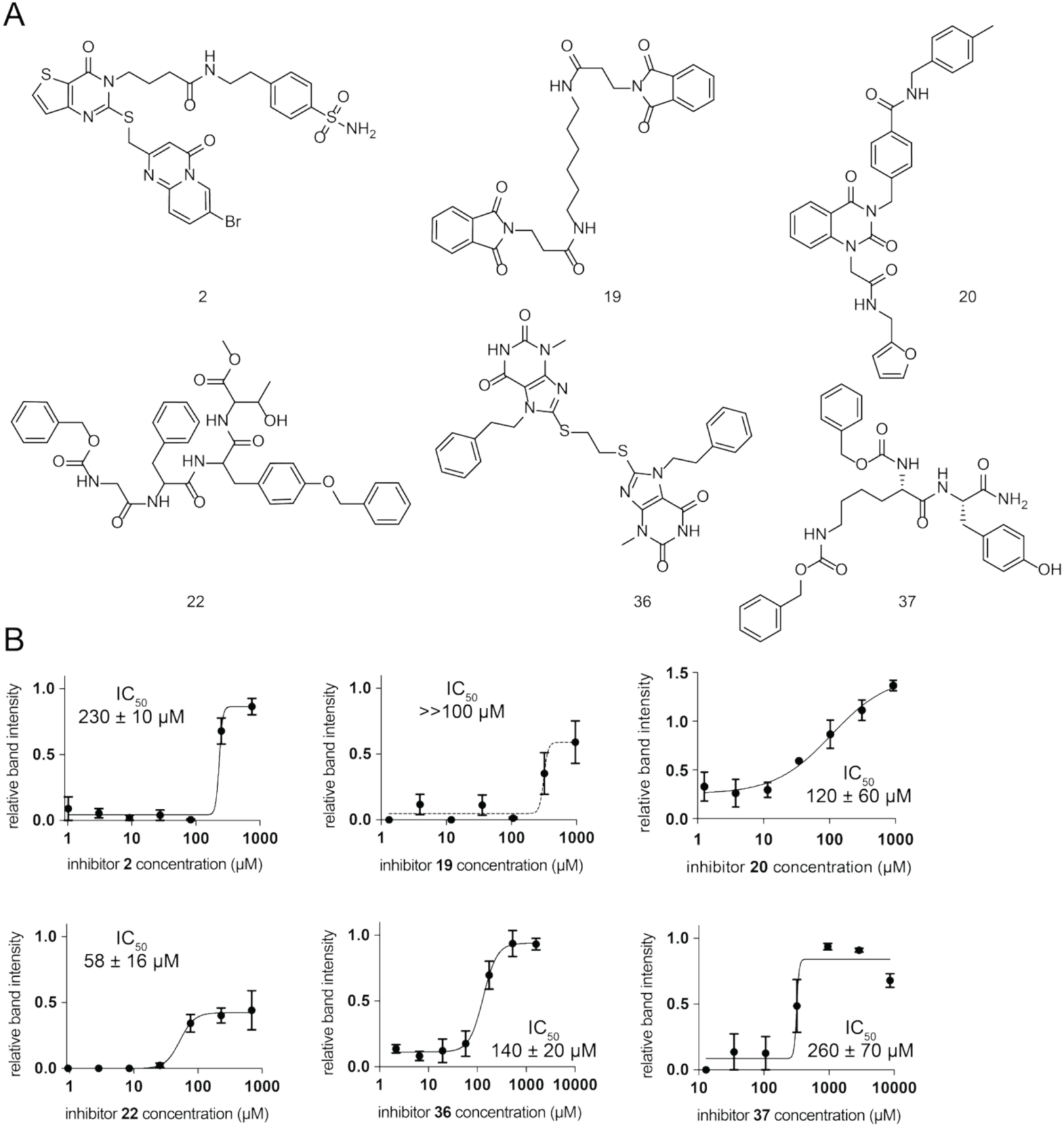
*Ll*Cap5 inhibitors. A) Structures of all biochemically validated inhibitors of *Ll*Cap5. B) Dose-response curves for *Ll*Cap5 inhibitors. Error bars represent standard error of the mean (S.E.M.) of a biological triplicate. IC_50_s are presented ± S.E.M. of biological triplicates. Curve fits for **19** did not afford confident IC_50_ values, but it is >>100 µM.

### *Ll*Cap5 inhibitors also inhibit a homologous Cap5 protein from a plant pathogen

After validating these inhibitors against *Ll*Cap5, we asked whether the inhibitors were specific to *Ll*Cap5 or if they could also target homologous Cap5 endonucleases in other bacteria. To address this, we focused on *Ps*Cap5 from the plant pathogen *Pseudomonas syringae*. We selected this homolog because it is activated by the same 3′,2′-cGAMP ligand,^32^ it may have implications for phage-based crop protection strategies, and it has recently been structurally characterized.^32, 33^ Comparison of the experimental protein structures of *Ps*Cap5 and *Ll*cap5 SAVED domains revealed that the proteins overlapped very well with a Cα RMSD of 1.4 Å (Fig. 3A). Both structures display conserved placement of the key residues that interact with the cGAMP ligand’s phosphodiesters (Arg234 and Ser274 in *Ll*Cap5, and Arg242 and Ser 277 in *Ps*Cap5, Fig. 3B).^31, 32^ These residues are crucial for the ligand recognition, as confirmed by alanine substitution mutations.^32^ Another essential residue, Arg281 in *Ll*Cap5 and Arg366 in *Ps*Cap5, recognizes the Hoogsteen edge of the guanine base of cGAMP in both structures, although the orientations of their side chains differ. Subtle differences in the ligand binding pockets are also evident. His138, located within a flexible loop in *Ps*Cap5, forms a hydrogen bond with phosphodiesters of cGAMP, whereas a corresponding loop in *Ll*Cap5 is disordered in the structure. Conversely, in *Ps*Cap5, Tyr304 contacts the *N*^2^ amino group of the guanine moiety of the ligand contributing to the specificity of ligand recognition (3′,2′-cGAMP vs 3′,2′-c-diAMP).^32^ However, in *Ll*Cap5, Tyr304 is replaced by Phe304, which results in the loss of direct recognition of the *N*^2^ group.

To assess whether the subtle differences between the pockets affected inhibitor activity, we re-screened *Ps*Cap5 with all 37 compounds previously identified as hits from the virtual screen. We observed both similarities and differences compared to the *Ll*Cap5 data. Notably, compound **4**, which was inactive against *Ll*Cap5, showed significant dose-dependent inhibition of *Ps*Cap5 (Fig. 3C–E). In fact, compound **4** is the most potent inhibitor of *Ps*Cap5 (IC_50_ = 11 ± 1 µM, Fig. 3E). Additionally, compound **20** showed ∼5⨉ higher potency against *Ps*Cap5 with IC_50_ = 24 ± 8 µM (Fig. 3F, compared to 120 µM against *Ll*Cap5). In contrast, compounds **2**, **19** and **22** did not noticeably inhibit *Ps*Cap5 (Fig. 3D), even though they inhibited *Ll*Cap5. Compounds **36** and **37** exhibit similarly low potency inhibition against both nucleases (Fig. 3G–H). Intrigued by the improved potency of compound **20**, we also tested ten analogs of it against *Ps*Cap5; however, none exhibited an IC_50_ lower than that of **20** (Table S1).

Since several SAVED domain binding inhibitors were effective against both Cap5 nucleases, we wanted to verify whether they interfere with the nuclease domains directly—potentially by sequestering active site ions or through other mechanisms—and thus might inhibit unrelated nucleases. Thus, we tested the most potent inhibitor of both *Ll*Cap5 and *Ps*Cap5 (compound **20**) against two unrelated restriction endonucleases we had on hand (Fig. S4). Compound **20** failed to inhibit both XhoI and DpnI, indicating that our inhibitors are not indiscriminate endonuclease inhibitors. Overall, some inhibitors selectively target *Ll*Cap5, others specifically inhibit *Ps*Cap5, and a few inhibit both cGAMP-activated Cap5 nucleases.

**Figure 3.**
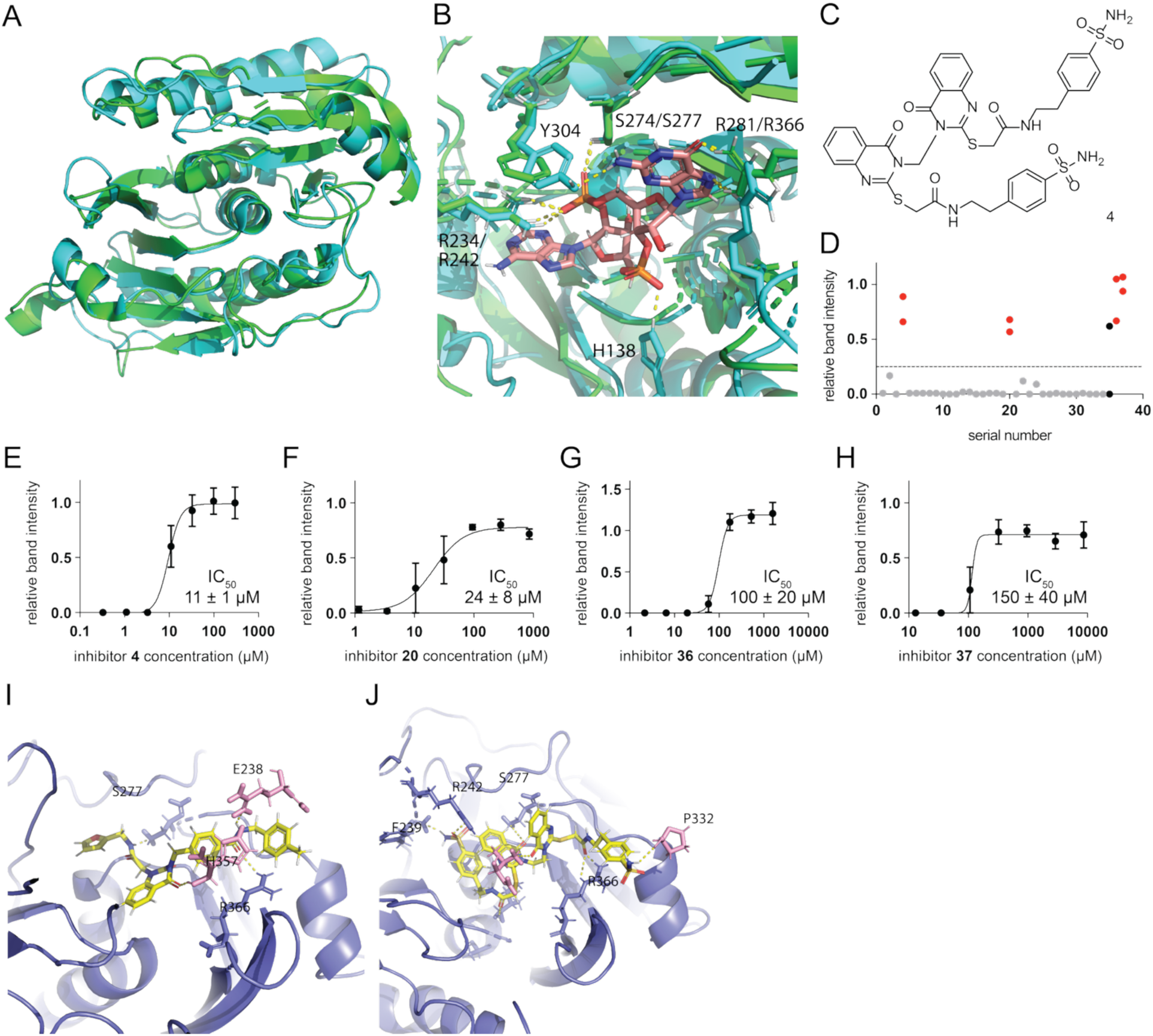
*Ps*Cap5 inhibitors. A) Superposition of the crystal structures of the SAVED domains of *Ll*Cap5^31^ in green and *Ps*Cap5^32^ in blue. B) Ligand-binding pockets of the superimposed structures from panel A. Critical residues Arg234 and Ser274 in *Ll*Cap5 (green) are conserved with residues Arg242 and Ser277 in *Ps*Cap5 (blue). Ligand is shown in pink. C) Structure of compound **4.** D) Rescreening of the compound library against *Ps*Cap5. Red dots represent four confirmed inhibitors. Grey dots represent compounds inactive in the initial test. Black dots are compounds that appeared active initially but were confirmed inactive upon retesting (second dot below threshold). Threshold at Y=0.25. E–H) Dose-response plots of inhibitors against *Ps*Cap5. Error bars represent S.E.M. of biological triplicates. IC_50_s are presented ± S.E.M. of biological triplicates. I–J) Docking results of compound **20** (I) and compound **4** (J) in *Ps*Cap5 ligand binding pocket, formed by the opposing surfaces of two SAVED domains of protomers A (purple) and B (pink). The inhibitors are shown in yellow.

### Docking suggests different protein interactions for *Ps*Cap5 inhibitors 4 and 20

To gain further insight, we examined the docking poses of compound **20** within the complete 3′,2′-cGAMP binding pocket formed by the two SAVED domains of each crisscross dimer in the *Ps*Cap5 tetramer.^32^ Five out of the top ten docking poses revealed hydrogen bond interactions with residues Ser277 and Arg366 on one side of the first SAVED domain (protomer A of the A/B crisscross dimer), which forms most of the binding pocket, as well as with Glu328 and His357 on the opposite side of the second SAVED domain (protomer B of the A/B crisscross dimer), that completes the ‘lid’ of the binding pocket via limited interactions (Fig. 3I). Notably, Ser277, Arg366, and His357 participate in 3′,2′-cGAMP binding in the crystal structure, suggesting that compound **20** may act as a competitive inhibitor interacting with both SAVED domains of the ligand-binding pocket.

In contrast, compound **4** exhibited a distinct docking profile that may account for its lower IC_50_, as illustrated in Figure 3J. Specifically, compound **4** forms hydrogen bonds with Ile219, Phe240, Arg242, and Ser277 of protomer A, as well as with Pro332 and Thr355 of protomer B. Notably, Arg242 and Ser277 participate in 3′,2′-cGAMP recognition.^32^ Together, these observations suggest that both compounds have potential as competitive inhibitors, though they likely interact with different key residues within the binding site.

### Compound 20 inhibits *Ps*Cap5 in cells

We aimed to access whether our inhibitors could enter the bacterial cells and effectively suppress Cap5 activity. For this, we focused on our two most potent inhibitors of *Ps*Cap5: compounds **20** and **4.** We incubated *E. coli* cells carrying a plasmid encoding *Ps*Cap5 and subsequently activated *Ps*Cap5 with the extrinsic addition of 3′,2′ -cGAMP, causing a severe restriction of bacterial growth (Fig. 4) consistent with the observations in a previous study.^32^ Thus, the number of viable bacteria was reduced at the 10^-1^ and 10^-2^-fold dilutions, and no viable bacteria were detected in the 10^−3^–10^-6^ dilution range. Because the expression of *Ps*Cap5 is leaky, the protein is produced even in the absence of IPTG.^32^ We then evaluated the in-cell efficacy of inhibitors by measuring their ability to rescue bacterial growth from cGAMP-Cap5-induced toxicity (Fig. 4). Upon the introduction of compound **20**, the population exhibited a remarkable recovery of almost 100-fold with the bacterial growth visible with 10^-3^-10^-4^ dilution range (Fig. 4B, C). These results strongly suggests that compound **20** can enter the cytosol of Gram-negative bacteria and inhibit the activation of *Ps*Cap5.

In contrast, compound **4** failed to rescue *E. coli* from cGAMP-Cap5-induced toxicity (Fig. 4C). We hypothesized that this failure was due to compound **4**’s inability to access the *E. coli* cytosol. To investigate this, we used the Entryway server^40^ to determine features in compound **4** that may be problematic. Indeed, compound **4** had less favorable features for accumulation in *E. coli*. Namely, while compound **20** has 9 rotatable bonds and a globularity value of 0.042, compound **4** has 17 rotatable bonds and a globularity of 0.125. Compounds with higher molecular flexibility (more rotatable bonds) and higher globularity are less likely to penetrate and accumulate within Gram-negative bacterial cells.^40^ Therefore, compound **20**’s enhanced ability to accumulate intracellularly likely explains its superior cellular activity despite similar *in vitro* IC_50_ values.

**Figure 4.**
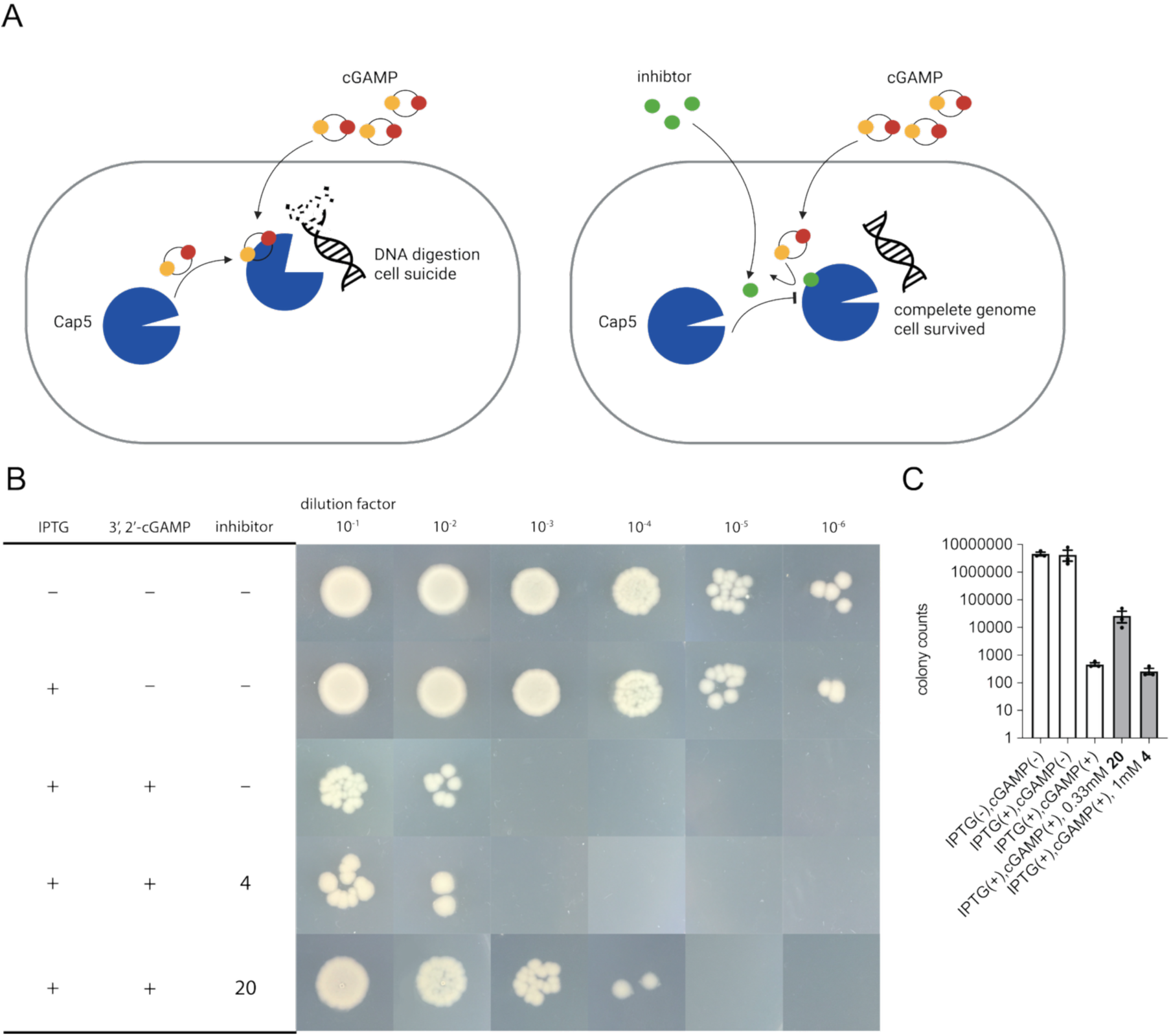
Inhibition of Cap5 in bacterial cells. A) Cartoon of the extrinsic activation of cellular Cap5 and bacterial growth inhibition. B) Representative images of serial dilution of *E. coli* culture carrying plasmid encoding *Ps*Cap5 after addition of activating cGAMP ligand. C). Bar plot of the recovery of viable cells with or without inhibitors **4** and **20** added. Error bars represent S.E.M. of a biological triplicate. Individual replicates are shown with circles.

### No virtual screening hits can activate *Ps*Cap5

A potential alternative outcome of synthetic ligands binding to the Cap5 ligand binding site is to activate the nuclease instead of inhibiting it. To explore this possibility, we considered that the 30 compounds which failed to inhibit either Cap5 in our biochemical assay might mimic the native ligand and activate Cap5. To test this, we re-screened the nuclease activity of *Ps*Cap5 in the presence of each of the 37 virtual screening ‘hits’, but this time in the absence of cGAMP. None of the compounds elicited noticeable nuclease activity (Fig. S5), suggesting that none could replicate the Cap5-activating function of cGAMP.

## DISCUSSION

In this study, we successfully identified the first chemical inhibitors of a CBASS defense system through a combination of virtual screening and biochemical validation. Our findings highlight the efficacy of docking-based strategies in identifying modulators of effector proteins, particularly those harboring defined ligand-binding pockets. As the structural landscape of phage defense systems continues to expand,^41^ the opportunity to leverage this strategy across different defenses grows increasingly viable.

We discovered compounds that inhibited the homologous Cap5 nucleases found in *L. lactis* and *P. syringae*. The compounds showed varied selectivity: some were more potent against *Ll*Cap5, others were more effective against *Ps*Cap5, and a few inhibited both enzymes with similar potency. This finding suggests that it is possible to either selectively target the phage defenses of individual bacteria or broadly target a class of related defenses, depending on the intended application.

Furthermore, we found that at least one of our inhibitors was able to permeate into the cytosol of Gram-negative bacteria—a requirement for most applications of inhibiting phage defenses. Notably, compound **20** targeted both *Ll*Cap5 and *Ps*Cap5 enzymes with the highest potency observed against *P*sCap5. Therefore, it may have application in sensitizing the plant pathogen *P. syringae* to phage treatments.

Despite multiple attempts, we failed to express a completely functional CD-NTase -Cap5 *Ll* or *Ps* CBASS defense operon in *E. coli* and observe clear anti-phage defense, which prevented us from validating the ability of our inhibitors to re-sensitize a host to phages. However, recent studies are beginning to demonstrate the feasibility of this approach.^17, 18^ Several studies have also explored the synergistic roles of small-molecule adjuvants in phage infection models.^19, 42^ A potential limitation of our study is the relatively low potency of the identified chemical inhibitors, which may explain why compound **20** did not fully restore viability of Cap5-expressing cells activated with the cGAMP ligand. More potent inhibitors may be required to effectively block phage defenses in wild-type bacteria. Alternatively, even partial inhibition of the defense may be sufficient to enable phages to escape the abortive infection process and destroy their targeted bacterial population.

Overall, this work establishes a foundation for targeting bacterial immune systems via chemical inhibition. It opens new avenues for studying the significance of anti-phage defenses in microbiomes and clinical settings and holds promise for enhancing the efficacy of phage therapy strategies against phage-resistant pathogens. Moving forward, future research can investigate the efficacy of these inhibitors in authentic phage-defense contexts and explore chemical modifications to enhance their potency and broad-spectrum activity across CBASS defense systems in other bacteria.

## METHODS

### Redocking cGAMP with *Ll*Cap5

The coordinates of the SAVED domain of *Ll*Cap5 protein with 3′,2′-cGAMP (PDB ID: 7RWS)^31^ was downloaded from the protein data bank and prepared in GOLD^34^ by adding hydrogens, deleting waters, and extracting the ligand. The extracted ligand was reset to its minimal energy conformation using the “clean up” function in Chem3D (Revity Signals Software, Inc). The cleaned-up ligand was then chosen as new ligand in GOLD for re-docking.

### High-throughput screening on supercomputer cluster

An 8 million lead-like 3D molecule structure library (including only molecules with molecular weight 300 to 800 Da) was downloaded from the ZINC 15 database.^43^ Openeye OMEGA FILTER^44^ was applied to remove PAINS molecules, then Openeye OMEGA QUACPAC was used to fix the protonation state at pH = 7.4. The virtual screening started with the preparation of the target protein *Ll*Cap5 (PDB ID: 7RWS).^31^ A docking .conf file was generated with GOLD v2022.2 and uploaded to a supercomputer cluster for virtual screening. For virtual screening, GOLD_Cluster_Computing v2022.1 was used with default settings. Following the initial screening, the top 70,000 molecules were picked for a higher accuracy docking, each docked with docking efficiency of 200% and 10 conformations per compound. From this second-round screening, the top 102 molecules were selected and further characterized via a biochemical assay.

### Identification of fingerprint, dimensionality reduction, and molecule clustering

Molecular structures were curated from an input file containing SMILES strings and parsed using RDKit 2025.03.3.^37^ RDKit Daylight-like fingerprints were computed using a fixed length of 2048 bits. All fingerprints were converted to NumPy^45^ arrays to enable downstream analysis. To visualize molecular similarity in low-dimensional space, principle components analysis (PCA) was applied to the binary fingerprint matrix using scikit-learn v1.6.0.^38^ Molecules were grouped using the K-Means algorithm in scikit-learn (KMeans, n_init=10, random_state=42, k=3) based on their PCA-transformed coordinates. To obtain chemically meaningful representatives for each cluster, a medoid selection strategy was implemented. For each cluster, the medoid molecule was defined as the compound with the minimal sum of Tanimoto distances to all other cluster members.

### Cloning, expression, and purification of recombinant proteins

The *Ll*Cap5 expression plasmid (codon-optimized for *E. coli*) on pRSF-1 was provided by Raven Huang.^31^ The *Ps*Cap5 expression plasmid (codon-optimized for *E. coli*) on pET28b(+) is the same as used previously.^32^ Both Cap5 proteins carry an N-terminal 6XHis tag followed by a SUMO tag. 100 mL *E. coli* BL21 (DE3) transformed with each expression plasmid was grown in LB with 50 µg/mL kanamycin, 220 rpm shaking, at 37 °C until OD_600_ reached 0.4−0.6. After cooling to 18 °C, expression was induced with 0.5 mM isopropyl-β-D-thiogalactopyranoside (IPTG), and cells were grown at 18 °C overnight.

The cultures were harvested by centrifugation, and the cell pellets were resuspended in lysis buffer (20 mM Tris-HCl, pH 8.0, 500 mM NaCl, 5% glycerol). Cells were lysed using a Fisher Scientific Sonicator FB505 on ice (55% amplitude, pulse time 2s/8s, total 5 min), followed by centrifugation at 15,000 rpm for 50 min at 4 °C to remove cell debris. The supernatant was loaded onto a 5 mL HisTrap^TM^ HP column (GE Healthcare). The proteins were washed using 25 mL washing buffer (20 mM Tris-HCl, pH 8.0, 500 mM NaCl, 30 mM imidazole, 5% glycerol) and eluted using elution buffer (20 mM Tris-HCl, pH 8.0, 500 mM NaCl, 500 mM imidazole, 5% glycerol). Peaks were detected with an AKTA Start system (GE Healthcare), and the fractions containing His-SUMO-tagged proteins were combined and cleaved with a His-tagged Ulp1 protease (Sigma SAE0067) in digestion buffer (20 mM Tris-HCl, pH 8.0, 150 mM NaCl, 5% glycerol) for 1 h at 30°C or overnight at 4°C. The reaction solution was loaded again onto the HisTrap column, and crude Cap5 protein was collected in the flow-through and washing buffer. The collected proteins were then concentrated with an Amicon® Ultra Centrifugal Filter, 30 kDa MWCO (UFC9030), and further purified on a HiLoad^TM^ 16/600 Superdex 200 pg size exclusion chromatography (GE Healthcare) column with reaction buffer (20 mM Tris-HCl, pH 7.4, 1 mM MgCl_2_, 0.5 mM MnCl_2_). Protein purity was assessed by SDS-PAGE with Coomassie staining. Samples were either frozen containing 10% glycerol or stored unfrozen with 40% glycerol at −20 °C.

### Inhibitor screening and dose-response DNA digestion assay

For all DNA degradation assays, BsaI-HFv2-linearized (New England Biolabs, R3733) *Ll*cap5 containing pRSF-1 plasmid (4,811 bp) was used as the DNA substrate. Assays were performed by incubating 50 nM *Ll*Cap5 or *Ps*Cap5 with 10 nM of the cyclic dinucleotide signal molecule 3′,2′-cGAMP (Enzo Life, BLG-c238-005) on ice for 10 min in Cap5 Reaction Buffer (20 mM Tris-HCl, pH 7.4, 1 mM MgCl_2_, 0.5 mM MnCl_2_) in a final reaction volume of 10 μL. The degradation reaction was initiated by the addition of 8 ng/μL DNA substrate, followed by incubation at 37°C for 15 min. Reactions were stopped by addition of 2 µL 6X Loading buffer (TriTrack, Thermo Scientific, R1161), and then 10 μL was separated on a 0.8% agarose gel (containing SYBR Safe stain, Invitrogen, S33102) in 0.5X TBE (0.045 M Tris-Borate, 1 mM EDTA). Gels were run at 100 V for 30 min and imaged with a Bio-Rad Universal Hood III gel scanner. The band was manually detected and normalized to positive control with no ligand added using Bio-Rad Image Lab 6.1. The activation assay was performed under the same experiment conditions without the addition of cGAMP.

### Restriction enzyme inhibition assay

The restriction enzyme inhibition assay was conducted using XhoI (NEB R0146S) and DPnI (NEB R0176S) in 1X rCutSmart Buffer (NEB B6004S). 1 µL 100 µM inhibitor **20** and 1 µL restriction enzyme were incubated on ice in 10 µL buffer for 10 mins. The degradation reaction was initiated by the addition of 8 ng/μL DNA substrate, followed by incubation at 37°C for 15 min. The post-reaction processing was conducted as described in the DNA digestion assay above.

### Docking mechanism study of compounds 4 and 20

The coordinates of the *Ps*Cap5 protein (PDB ID: 8FMH)^32^ were downloaded from Protein Data Bank and prepared for docking by adding hydrogens, deleting waters, and extracting the ligand using GOLD. The docking efficiency was 200%, and 20 conformations were generated for each molecule.

### In cell inhibitory activity test of compounds 4 and 20

The same *E. coli* BL21 (DE3) cells carrying the *Ps*Cap5 expression plasmid from above were grown in LB medium with 50 μg/mL kanamycin at 37 °C with 220 rpm shaking to an OD_600_ of ∼0.6 (1 cm pathlength). Then, 200 µL of the culture was transferred to multiple wells of a 96 well plate and incubated at 37°C for 3 h with or without 0.1 μM IPTG and/or 0.2 mM 3′,2′-cGAMP and 0.33 mM compound **20** or 1 mM compound **4**. The bacterial cultures were then serially diluted from 10^-1^ to 10^−6^, and 5 μL of each was spotted onto LB agar petri dishes supplemented with kanamycin. After overnight incubation, the plates were photographed and colonies formed by viable cells were counted manually.

## AUTHOR CONTRIBUTIONS

Conceptualization, C.Z., J.P.G.; Methodology, C.Z., J.P.G.; Investigation, C.Z., O.R.; Writing – Original Draft, C.Z.; Writing – Review & Editing, C.Z., O.R., J.P.G.; Visualization – C.Z., J.P.G.; Supervision, J.P.G.; Funding Acquisition, J.P.G.

## MATERIALS & CORRESPONDENCE

Correspondence and material requests should be addressed to J. P. Gerdt.

## ACKNOWLEDGMENTS

We thank Raven Huang for providing the *Ll*Cap5 plasmids. The research was supported by a National Science Foundation CAREER award (IOS-2143636) to J.P.G., and a Camille Dreyfus Teacher-Scholar Award (TC-24-028) to J.P.G. This research was supported in part by Lilly Endowment, Inc., through its support for the Indiana University Pervasive Technology Institute.

## SUPPORTING INFORMATION

**Figure S1.**
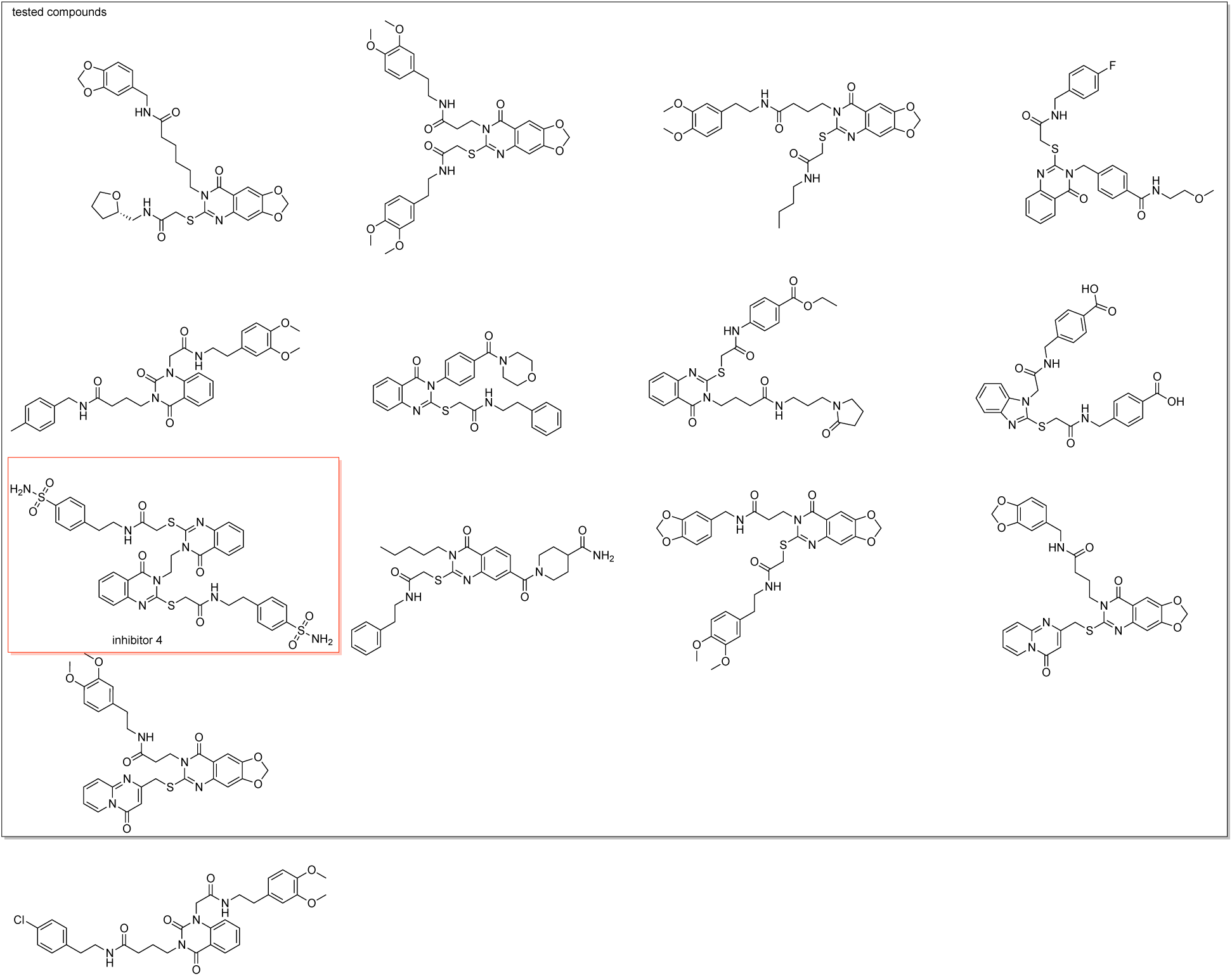
Group A of the top screening results. The boxed ones were purchased for inhibitor screening, the highlighted ones were validated as inhibitors.

**Figure S2.**
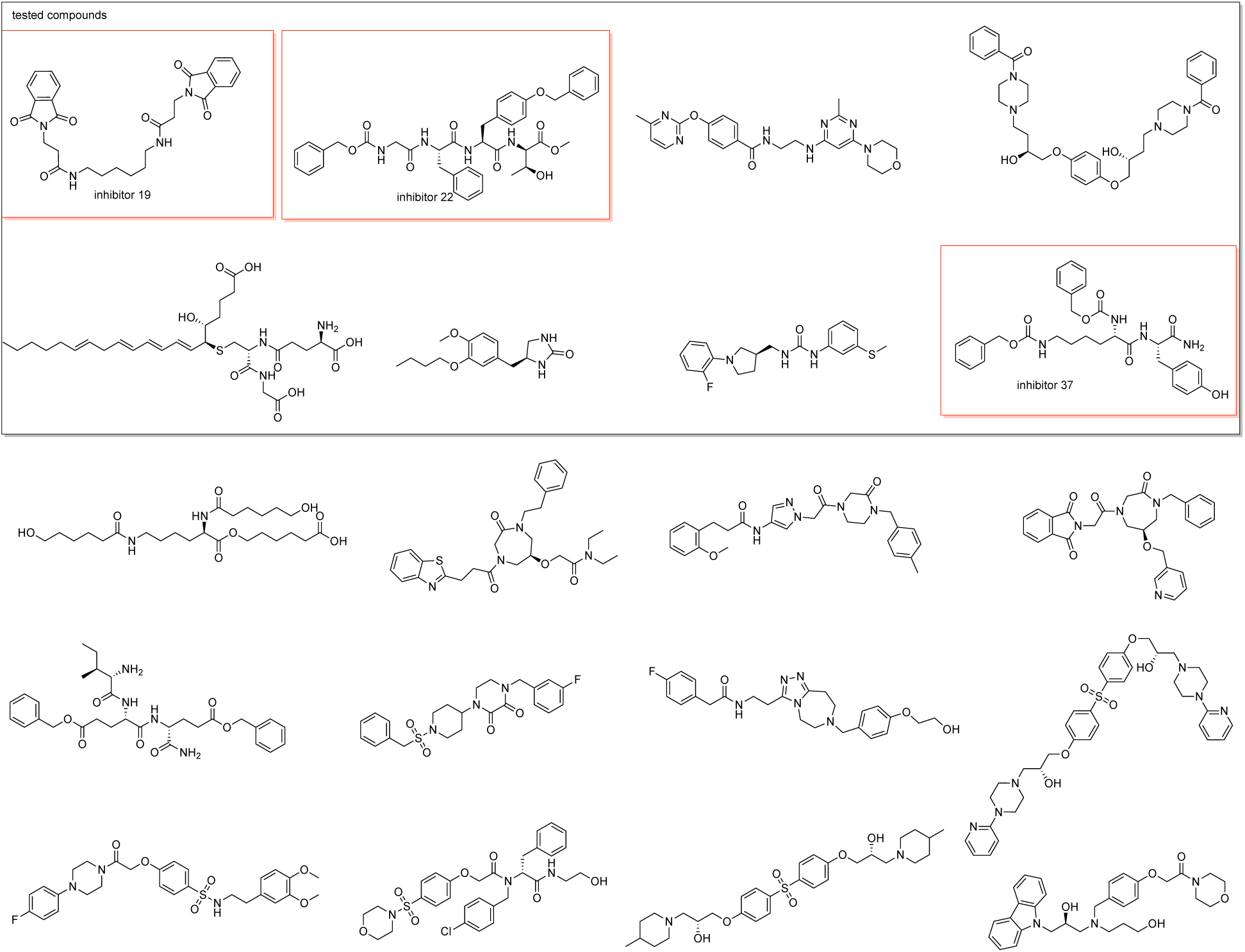
Group B of the top screening results. The boxed ones were purchased for inhibitor screening, the highlighted ones were validated as inhibitors.

**Figure S3.**
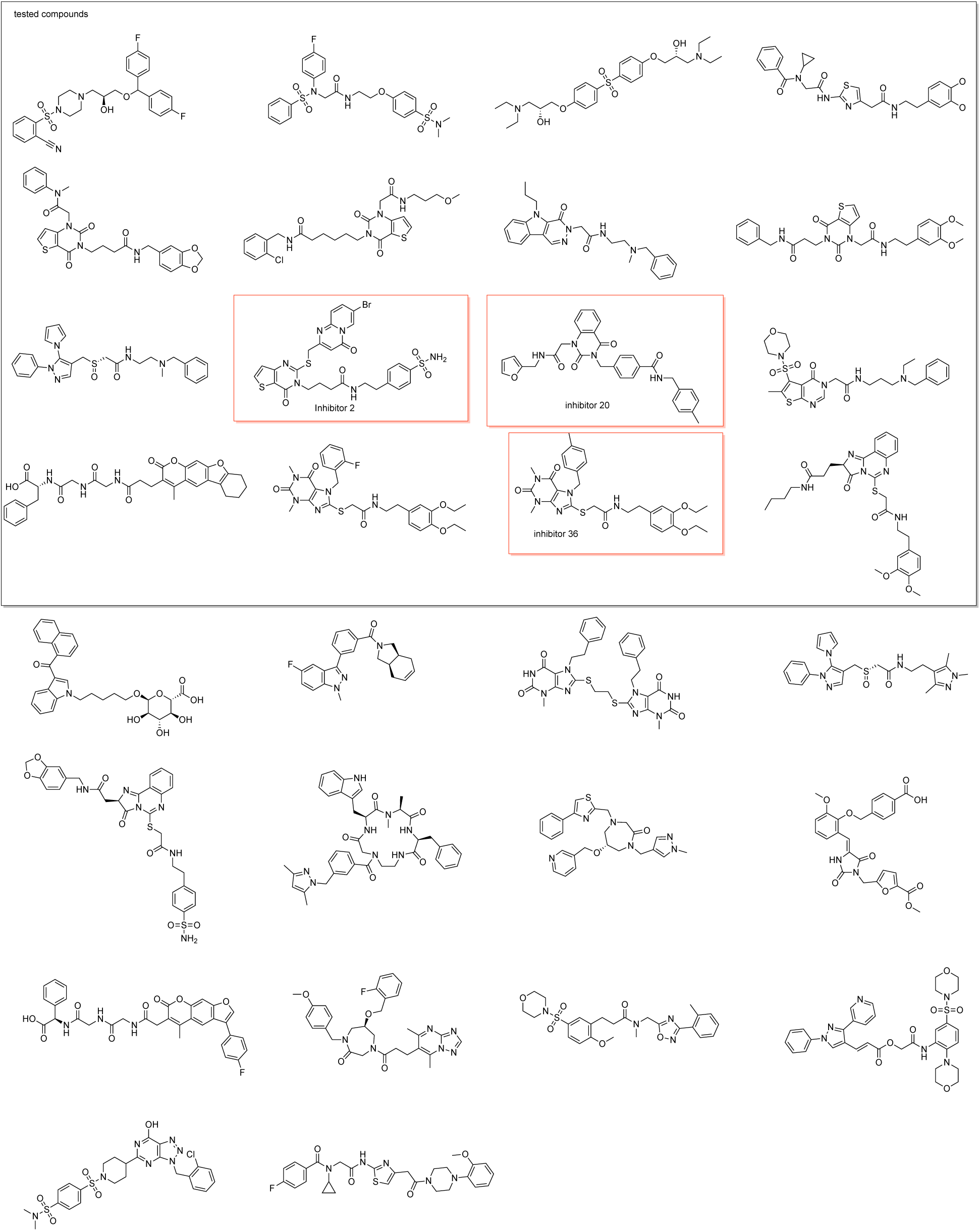
Group C of the top screening results. The boxed ones were purchased for inhibitor screening, the highlighted ones were validated as inhibitors.

**Figure S4.**
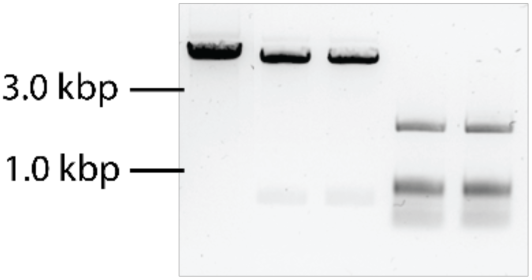
Test of general endonuclease inhibition. Lane 1: Positive control, linearized plasmid. Lanes 2&3: Xhol with and without inhibitor **20.** Lanes 4&5: DpnI with and without inhibitor **20.**

**Figure S5.**
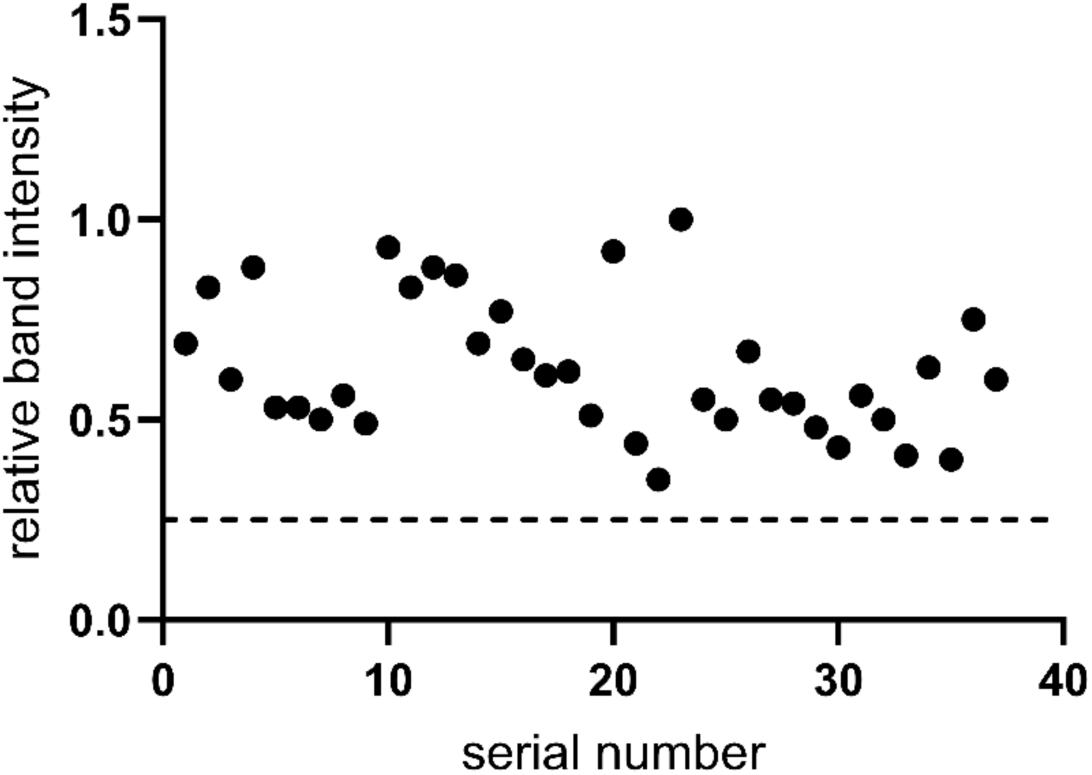
Screen for PsCap5 agonists. No molecules induced substantial DNase activity in the absence of cGAMP.

**Table S1.**
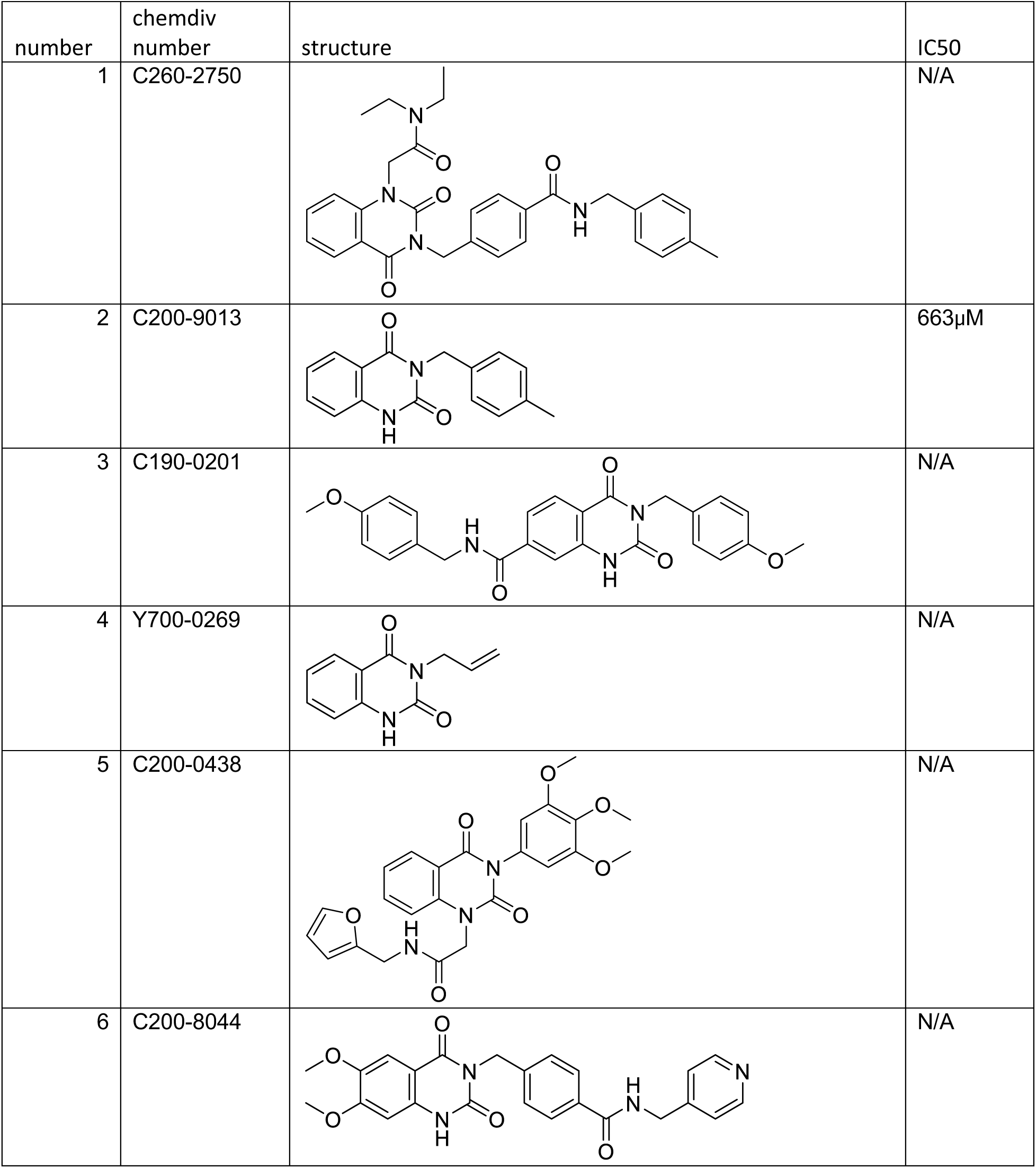
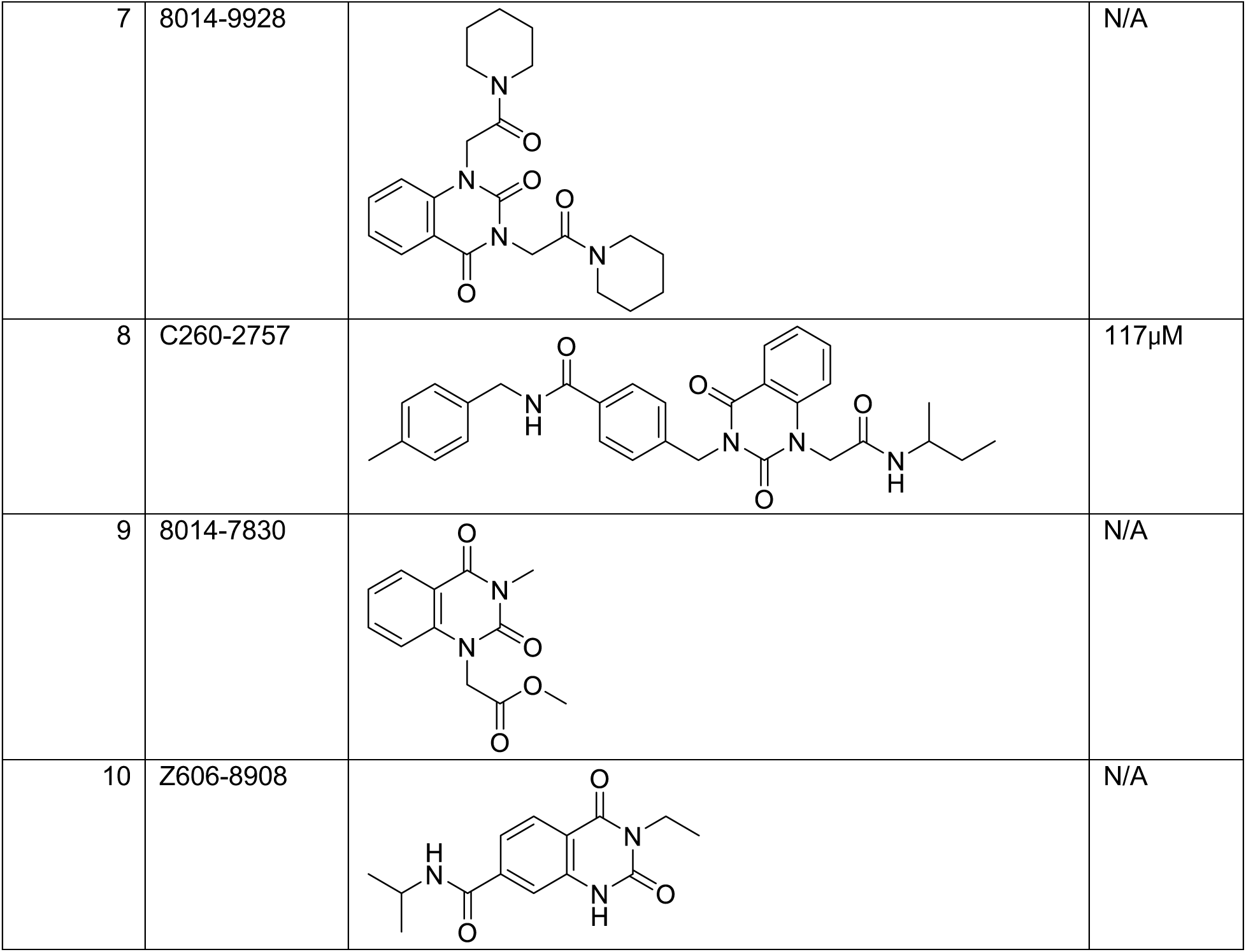
IC_50_s of 10 analogs of compound 20.

| number | chemdiv number | structure | IC<sub>50</sub> |
| --- | --- | --- | --- |
| 1 | C260-2750 |  | N/A |
| 2 | C200-9013 |  | 663μM |
| 3 | C190-0201 |  | N/A |
| 4 | Y700-0269 |  | N/A |
| 5 | C200-0438 |  | N/A |
| 6 | C200-8044 |  | N/A |
| 7 | 8014-9928 |  | N/A |
| 8 | C260-2757 |  | 117μM |
| 9 | 8014-7830 |  | N/A |
| 10 | Z606-8908 |  | N/A |

